# Natural selection subsumes and unites multiple theories of perceptual optimization

**DOI:** 10.1101/507459

**Authors:** Victor Quintanar-Zilinskas

## Abstract

Perceptual processing is thought to occur in two stages^1,2^: compression of the environment’s information, followed by inference of the environment’s state. While inference is believed to be Bayes-optimal^3–5^, the model for optimal compression is comparatively less well-established: multiple signal processing frameworks have been used to explain empirical outcomes^1,2,6–8^ and hypothesized to align with Darwinian fitness^1,2,7–9^. Here I endeavor to constrain the model space by framing fitness as a function of the compression scheme and optimizing fitness directly. I find that signal processing and fitness objectives often deviate; for example, in a finite game where risk-avoidant bet-hedging^10^ beats maximization of average performance. However, when average performance determines fitness, then both are maximized by minimized cost-weighted signal distortion. Meanwhile, the signal processing metrics of estimation error and mutual information align with these latter objectives only under very specific circumstances. Taken together, my results establish a Darwinian (nested) hierarchy of perceptual compression objectives, which extends the existing understanding of optimal compression by incorporating not just signal processing but also economic frameworks, and reconciles the extant usages of the former by clarifying their respective domains of applicability. The cost-weighted distortion objective’s evolutionary importance is additionally intriguing because this objective is potentially congruent with a broad set of experimental findings.

---

To evaluate the fitness of perceptual compression schemes, I employ a framework in which an organism: is presented with n items, perceives them through its perceptual interface^11^, chooses one, and adds the item’s utility value to its total lifetime utility score. Not all interfaces are equally fit: their fitness hinges on their effective mapping of high- and low-utility items to different percepts, thereby enabling low-utility choices to be avoided. Also, the interface represents a compression: multiple items are mapped to the same percept, possible real-world items outnumber the possible percepts, and the mapping does not fully preserve the details of an item’s real-world attributes (Fig. 1A). Notably, this framework’s discretization of choice options renders the compression amenable to formal mathematical parameterization, while its discretization of choice instances and utilities renders the compression’s fitness consequences measureable. Despite these abstractions, the framework retains perception’s fundamental relationship to fitness: the information it transmits to the organism informs actions^12^.

**Figure 1.**
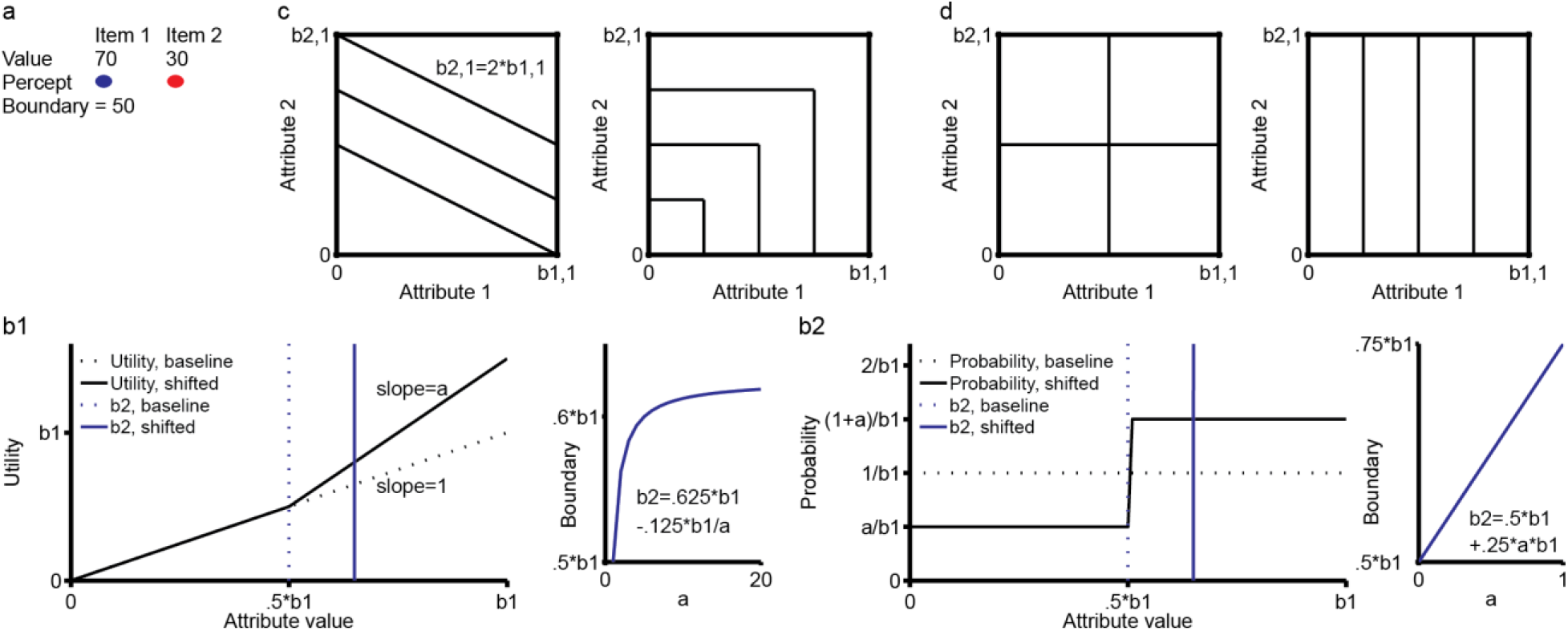
Illustration of utility-maximizing assignment of real-world items to percepts. **a**, Individual real-world items (1 and 2) have quantifiable attributes. Perceptual interfaces inevitably compress this information; here, an interface with a 1-bit output perceives items as blue or red, based on one particular attribute value being above or below a boundary. The organism favors blue options when choosing an item. **b**, Each pair of plots illustrates how (for a 1-bit interface making binary decisions) a particular deviation from a simple baseline utility landscape shifts the optimal perceptual boundary. **c**, The boxes in this panel (and the next) show the optimal division, by a 2-bit interface that makes binary decisions, of an item landscape in which utilities are determined by two uncorrelated attributes. Right and left boxes respectively segment items whose utilities exhibit situation-specific single-attribute dependency or are determined by their attribute sum. These interfaces are free from item-percept assignment constraints. **d**, These landscape divisions are optimal subject to the constraint that attributes are perceived separately. The right-box division is “optimal” (e.g., in environment 2) when utility hinges on attribute 1 far more than on attribute 2. The left-box division is favored evolutionarily when utility is similarly dependent on both attributes, or even in versions of environment 2 (where b_2,1_ ≫ b_1,1_) in which this “risk averse” division benefits from avoidance of rare, but large, utility-harvesting failures.

In the present study, I apply the above framework across several decision-making environments, each characterized by a set of selection pressures and distribution of item utilities. In each environment, certain interface parameters, cast as alleles that are subject to selection, are shown to be more adaptive than others, and perhaps even optimal in that particular environment. The various congruencies revealed between selected-for parameterizations and well-known compression and decision strategies collectively provide an expanded understanding of perceptual processing in the natural world.

In environment one, fitness is linearly proportional to average decision utility, item utilities are a function of a subset of their real-world attribute values, and the values of each utility-relevant attribute are contained within the interval [0,b_a,1_], where a is the index of a specific attribute. In a version of this environment simplified even further, utility depends linearly on just one attribute (in which case, I write b_a,1_ as b_1_), and item probability is uniform over the attribute space. In such a one-dimensional utility landscape, it is adaptive to associate percept boundaries with specific values of the utility-determining attribute, and assign all items between boundaries b_i_ and b_i+1_ to the same percept. Thus, an interface with β output bits divides the item space into m = 2^β^ percepts P_i_ (i = 1:m), whose index indicates both the percept’s ordinal desirability (relative to other percepts) and its upper boundary b_i_. While b_1_ is a constant, the other b_i_ (i=2:m) can be regarded as the alleles subject to selection.

In this environment, there is congruence between the utility maximization objective and the minimization of signal distortion, with the loss function being the compression’s utility cost (details in Supplementary Equations, Part 1). These objectives are demonstrably equivalent in the general case of a β-bit interface and n item choices (Supplementary Equations, Part 2). They are also equivalent, when β = 1 and n = 2, on utility landscapes exhibiting simple deviations from the assumptions of 1) a linear attribute-utility relationship 2) uniformly probable item attribute values (Fig. 1b1 and 1b2; Supplementary Equations, Parts 3 and 4; respectively). Moreover, these results straightforwardly extend to multidimensional utility landscapes. The extension is trivial for an organism with an idealized interface^11^ that can map items to percepts with infinite flexibility. If utilities are unitary (e.g. a weighted sum of attribute values), then the interface will in effect rearrange the items onto a unidimensional utility continuum (Fig. 1c, left).

Alternatively, if the organism experiences a variety of situations, each corresponding to a different utility-determinative attribute, then each dimension individually can be optimally divided (Fig. 1c, right). Granted, real-world interfaces are often composites whose individual components (e.g., vision and olfaction) are constrained to processing a subset of item attributes (e.g., reflected wavelengths and emitted chemical compounds). Under such circumstances, the utility and distortion loss objectives will dictate equivalent segmentations of individual dimensions and, at least for a two-bit interface making binary decisions between items from a two-dimensional utility space (Fig. 1d; Supplementary Equations, Part 5), an equivalent assignment of interface bits to the available dimensions.

Notably, when I set β = 1 and n > 2, the utility and distortion loss objectives demonstrably (Supplementary Equations, Part 6) diverge from the frequently-studied objectives of mutual information maximization and utility estimation error minimization^13^. The latter two are favored by selection with respect to neither items perceived nor items chosen; stated alternatively, selection favors neither the optimization of perceptual accuracy nor of action outcome prediction. Still, optimized action outcome prediction has been previously shown to provide a lower bound on utility obtained.

Environment two is mostly similar to environment one; the main differences, seemingly subtle, are that fitness is determined by utility accumulated over a finite lifespan (duration measured in decisions) rather than by average decision utility, and that after every generation, the fixed-size population undergoes a membership refresh during which the lowest-utility members leave no offspring. Additional key features of this environment: individual decisions depend on just one attribute, the utility landscape is two-dimensional, and although b_1,1_ ≪ b_2,1_, decisions contingent on attribute 2 are so rare that mean utility maximization dictates the allocation of both perceptual bits to attribute 1. Organisms interact with this environment via a 2-bit interface configured to either maximize mean utility (“optimal”, Fig. 1d, right) or avoid large utility-harvesting failures during those few decisions contingent on attribute 2 (“risk averse”, Fig. 1d, left).

Simulations of competition between these interface strategy alleles (details in Methods) mostly favor “optimal”; notably, all observed fixations of “risk averse” occur when reproduction is restricted to the best-performing 20%, and this outcome is also favored by short organism lifetimes (Table 1). Thus, the victories of “risk averse” can best be understood as successful bet-hedges^10,14^, defined as the sacrifice of maximized average performance in order to avoid unacceptable risks (such as falling into the population’s non-reproductive 80%), or as demonstrations of the advantageousness of risk-taking in competitions that reward relative standing^15,16^. These results show that when reproductive fitness deviates from linear dependence on average decision utility, strong correspondence between distortion loss and fitness objectives is still possible, but cannot be assumed.

**Table 1.** Simulations of competition between “risk averse” and “optimal” interface alleles

| Condition | Lifetime Decisions | Probability of high-stakes (HS) scenario | Risk averse allele, initial prevalence | Instances of “risk averse” allele fixation |
| --- | --- | --- | --- | --- |
| A | 1000 | .02 | 80% | 8 |
| B | 1000 | .015 | 80% | 0 |
| C | 100 | .015 | 80% | 7 |
| D | 100 | .015 | 50% | 1 |
| E | 100 | .02 | 50% | 9 |
| F | 100 | .02 | 20% | 1 |
| Comparison | P<sup>1</sup> | Parameter |  |  |
| A vs. B | .0007 | p(HS) |  |  |
| B vs. C | .0031 | Lifetime length |  |  |
| C vs. D | .0198 | Initial prevalence |  |  |
| D vs. E | .0011 | p(HS) |  |  |
| E vs. F | .0011 | Initial prevalence |  |  |

<sup>1</sup>Fisher’s exact test, two-sided

In environments three through six, item utilities are not fixed in evolutionary memory; instead, they must be experienced and learned. It is assumed that percept utilities are learned, and that percept preference order is established, rapidly. It is further assumed, except when otherwise noted, that average post-learning decision utility determines fitness. Environments three and four consist of 100 items with randomly sampled utilities; the expected value derived from a particular item-percept assignment is computed as described in Methods and Fig. 2.

**Figure 2.**
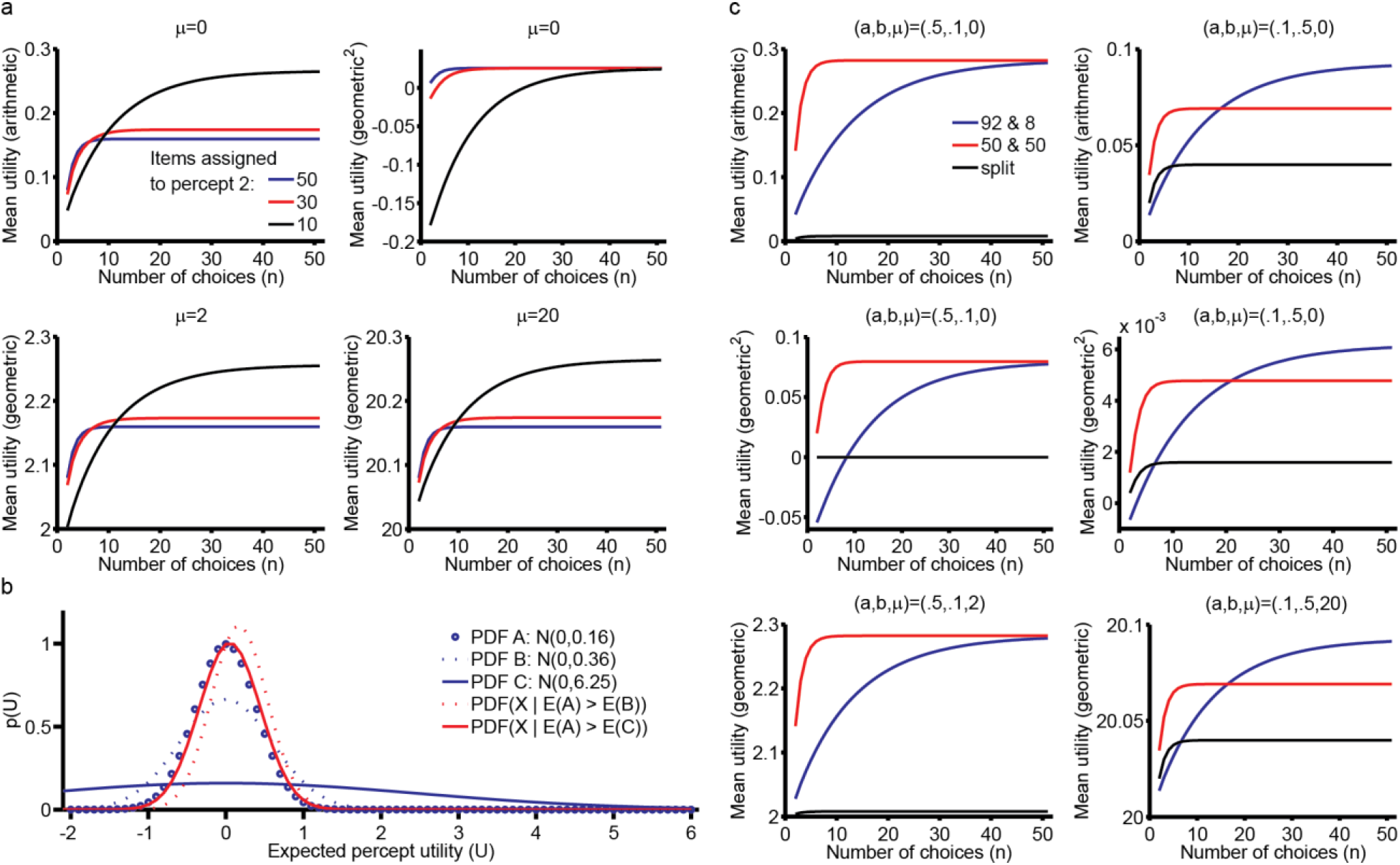
Optimizing the assignment of unknown-utility items to percepts whose utilities are learnable. **a**, Displays mean (arithmetic and geometric) decision utilities obtained using a 1-bit interface to distinguish amongst 100 items with utilities drawn independently from N(μ,4). **b**, When percept variances are similar (A vs. B), the low-variance percept’s high-utility random values are more likely than its low-utility random values to become learned preferences (dotted red line). However, this likelihood distinction mostly disappears (solid red line) when percept variances differ significantly (A vs. C), because the high-variance percept’s value almost wholly determines which percept is preferred. **c**, Here, item utilities belong to one of two clusters, μ+x_1_+N(0,b^2^) and μ+x_2_+N(0,b^2^); x_1_ and x_2_ are themselves drawn from N(0,a^2^). Line labels indicate either the sizes of percept-assigned clusters, or that each cluster’s members are “split” between percepts.

Fitness in environment three is determined by the interface’s extraction of value from uncorrelated, randomly-valued items. The top plots of Fig. 2a reveal (for n > 9) that in this endeavor there is a trade-off between pursuing large gains (black line) and steady gains (blue line). (The bottom plots show that if mean item utility is high, then reward is reliably available and the trade-off is mitigated or even eliminated.) Fig. 2b illustrates the genesis of this trade-off; it clarifies why the pursuit of large gain potential from one percept comes at the cost of decreased gain potential from the other, and in turn suggests that unbalanced percepts deliver low geometric mean utility because their pursuit of utility is undiversified^17^. The ultimate advantageousness of these alternative reward pursuit strategies depends on whether the environment selects for arithmetic or geometric mean utility maximization; thus, as in environment two, bet-hedging perception is potentially favored over utility-maximizing perception.

In environment four, meanwhile, item utilities are correlated: specifically, they belong to one of two clusters. I find (Fig. 2c) that even when within-cluster utility variation appreciably exceeds between-cluster variation, assigning the clusters to distinct percepts (red lines) delivers greater mean utility, arithmetic and geometric, than does splitting each cluster’s items evenly between the percepts (black lines). The latter strategy is potentially advantageous only if items from one of the clusters are very rarely encountered (blue lines), in which case environment four approximates environment three.

Environment five’s similarities to the simplest version of the first are that one attribute determines utility and that utility declines linearly from the peak-utility attribute value; what differs is that the utility peak, instead of being fixed, now assumes a random value within the interval [0,b_1_], or else within [−π,π] if the attribute space is radial rather than linear. Notably, the randomization of the utility peak can bring the estimation error objective into alignment with the utility and distortion loss objectives. More specifically, the three objectives align in the linear attribute space when β = 1 or n = 2 and in the radial attribute space for general β and n (Supplementary Equations, Part 7), and remain aligned in the radial attribute space (if n = 2) even when the probability distribution of items across the utility space is non-uniform (Supplementary Equations, Part 8).

The significance of these results comes into focus upon considering a more biologically realistic formulation of environment five (and four), in which the organism encounters a multiplicity of decision situations, each with a unique utility-maximizing attribute value (or, unique item cluster utilities). If environment one demonstrated that natural selection favors the selective recognition of an environment’s highest-utility items, then these results demonstrate that natural selection favors estimation accuracy across all items when all items have the potential to (sometimes) be the environment’s highest-utility item. On that note (Supplementary Equations, Part 8), efficient coding of the attribute space may sometimes be more adaptive than efficient coding of the environment’s item distribution; just because certain items are common doesn’t mean that differentiating amongst them is particularly important to organismal performance.

The principle that unites all of the present study’s findings is that natural selection favors perceptual interfaces revelatory of utility gradients that inform decisions. My formalization of this seemingly simple notion, which has enjoyed decades^7,18,19^ of acceptance, advances understanding in several important and unanticipated ways. First, it differentiates it from the notion^7,8^ that selection maximizes total utility harvesting, and shows that instead, interfaces calibrated to maximize total decision utility are a subset of all gradient-revelatory interfaces potentially favored by natural selection. Environments two and three, for example, showcase the adaptiveness of interfaces calibrated to avoid an environment’s lowest-utility outcomes (and arguably, by propagating interfaces specialized for the processing of risk-indicative stimuli^20,21^, so too does nature). Second, it differentiates it from the notion, also enjoying decades of acceptance^1,3,22^ (despite being incorrect), that selection favors maximally-informative (i.e. error-minimizing) interfaces, and instead shows that these are adaptive only to the extent that they abet the exploitation of utility gradients. This latter result further clarifies that maximally-informative interfaces are favorable under decision conditions similar to those faced by task-general interfaces that process low-level stimulus features^1^. Third, it presents cost-weighted distortion minimization as a broadly useful framework that can account for perceptual adaptation to several types of environmental variation, such as task structure (e.g. selective identification of one vs. multiple peak-utility items), utility gradients (e.g. Fig 1b1), and environmental item probabilities (e.g. Supplementary Equations, Part 8). Though this framework’s experiment interpretation utility is already established^6–8^, future studies may bridge it with an expanded set of empirical findings, such as categorical color perception^23,24^ and the effects of reward-associated plasticity on sensory cortex stimulus tuning^25,26^; after all, that plasticity is induced by previously-unencoded environmental utility gradients.

## Methods

### Interface allele competition

Three populations are simulated in parallel. The size of each is fixed at 100 individuals. Individuals differ according to whether their perceptual interface is “risk averse” (RA) or “optimal” (O). At simulation initialization, the prevalence of O is either 20%, 50%, or 80%; all three populations are initialized equivalently. Then, within each population, a lifetime of decisions is simulated for each member (the nth member of each population faces the same life decisions), the members are ranked by their lifetime’s accrued utility, and the membership of the next generation is generated according to rules specific to each population. Lifetime simulation and population refresh procedures repeat until each population converges to fixation for one of the two alleles.

The first and second sets of refresh rules are simple: the bottom 80% and 50% (respectively) of the utility accumulators are eliminated, and the set remaining is copied until the population size is restored to 100. The third set eliminates the bottom 10% and replaces each member thereof with an individual whose probability of exhibiting each allele is proportional to the allele’s prevalence in the top 90%.

The decisions made within a lifetime involve choosing between two items sampled (with uniform probability) from the utility landscape in Fig. 1d, with b_1,1_ = 100 and b_2,1_ = 1100. Most decision utilities are contingent upon attribute 1; those contingent upon attribute 2 are deemed “high stakes” (HS), and arise with probability p(HS). In these simulations, p(HS) = .02 or .015; the result in Supplementary Equations, Part 5 indicates that the O interface (Fig. 1d, right) achieves higher average decision utility than the RA interface (Fig. 1d, left) when p(HS) < .022.

### Statistics

In Table 1, the parameterization conditions A-F were each simulated 10 times. Then, the “risk averse” fixation probability was compared (Fisher’s exact test, two-sided, n=20) across various pairs of conditions differentiated by just one parameter.

### Expected utility computation

Because percepts are composed of items whose utilities are drawn from normal distributions, the distributions of mean percept values are also normal. Also, once percept utilities are learned, the organism will preferentially select items mapped to higher-utility percepts; a percept’s expected value, therefore, is related to its position in the percept preference order. If it is further specified that the interface has N percepts, and that a particular sampling of item utilities results in percept 1 being the highest-utility percept, then the expected value of percept 1, E(P_1_), can be expressed as

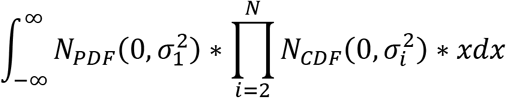

Alternatively, if E(P_1_) < E(P_2_) and E(P_1_) > E(P_i_) ∀ i > 2, then

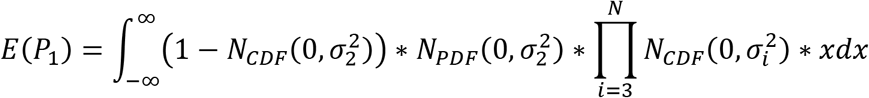

The expressions for all expected percept values across all preference orderings are constructed similarly. In the present study, these expressions are evaluated by numerical integration. The results in Fig. 2a and 2c are obtained by integrating over [−5,5] with dx = .001, those in Fig. 2b are obtained by integrating over [−6,6] with dx = .01.

A notable feature of the above expressions is that each percept has an associated σ-value. These are computed individually for every percept across all item-percept assignments, using standard methods. For example, for a 25-item percept with utilities drawn from N(0,1), σ = 1/5.

Upon computing the ordering-specific expected percept values, it becomes possible to combine them to compute the mean utility obtained by a particular assignment of 100 items to the available percepts. Specifically, the respective arithmetic and geometric mean utilities are (E(O_1_)+E(O_2_))/2 and (E(O_1_)*E(O_2_))^1/2^ (in Fig. 2, when some E(O_2_)*E(O_2_) < 0, I report the square of the geometric mean), where the O_j_ represent different percept preference orderings. Letting O_j_ denote the ordering in which j is the preferred percept, defining E(P_j,i_) as the expected value of percept i in ordering j, and defining S_i_ as the size (in items) of percept i,

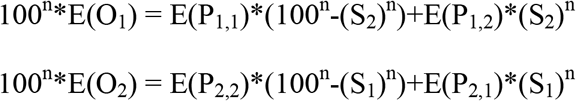

## Author Information

V.Q.-Z. declares no competing interests. Correspondence and requests for materials should be addressed to V.Q.-Z.

**Code and Data** are accessible at github.com/victorqz1/PerceptualCompression1.

## Supplementary Equations

**Note**: in these analyses, the range of utility values is assumed to be spanned by a density of items sufficient to justify an integral approximation for the calculation of: utility estimation error, item-percept mutual information, expected utility, and distortion loss.

**Part 1**: derivation of the objective function for distortion loss minimization

Distortion during a single decision: given a set of n items available during a particular decision round, one can label their utilities x_1_:x_n_, indexed in order of descending utility. A perceptual system that transmits environmental information perfectly would obtain utility x_1_, as would a perceptual system that assigns the x_1_-utility item to a category of its own. However, if there are k items that appear to be of the same, most-preferred percept, then the expected utility of the decision round is 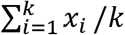. The distortion loss is the expected magnitude of the utility loss that results from the perceptual system’s compression of the world’s information; it’s magnitude is 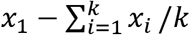.

Total distortion (TD), then, is simply the sum of the distortion losses over all possible utility value sets X = {x_1_,…,x_n_} when the interface uses the boundaries B = {b_2_,…,b_m_}. As an illustration of how boundaries affect the distortion, consider the set Q of all item sets {x_1_,…,x_n_} that satisfy b_i_ < x_1_:x_k_ < b_i+1_ < x_k+2_:x_n_. The contribution of Q to TD is

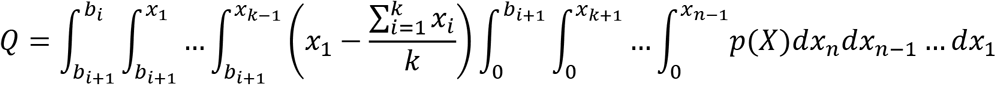

Note that the limits of integration reflect the ordinal arrangement of the x_i_, and their position relative to the b_i_. Note also that the distortion depends on p(X). Generalizing this expression across all boundaries i and best-percept item counts k, and noting that Q_i1_ = 0, yields 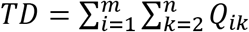.

**Part 2**: proof that for the general game with m percepts, n choices, a linearly increasing and unidimensional utility continuum, and a uniform distribution of item probabilities over that continuum, the perceptual category boundaries that optimize total distortion (TD) and utility (U) are equivalent

Approach: the optima of U and TD are equivalent if ∀ i, dU/db_i_ = C*dTD/db_i_, C constant.

**2.1**: expression for dU/db_i_

Recall from main text that: b_i_ and b_i+1_ are the upper and lower boundaries of P_i_; note also that b_m+1_ = 0, E(P_i_) = (b_i_+b_i+1_)/2, and

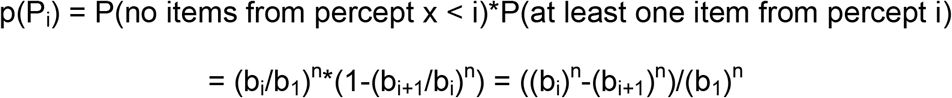

Thus, utility can be written as

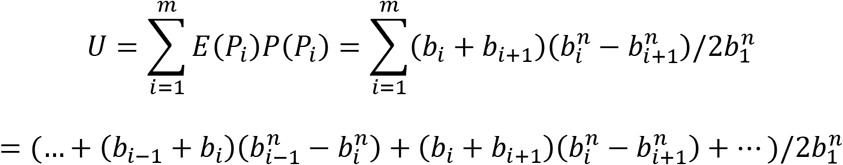

Setting dU/db_i_ = 0 and solving yields

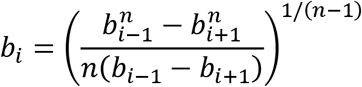

**2.2**: simplifying Q_ik_

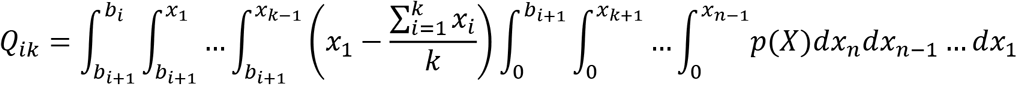

First, set aside the p(X) term; in this environment, it is constant ∀ X Second,

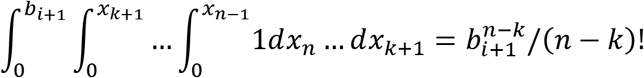

Third,

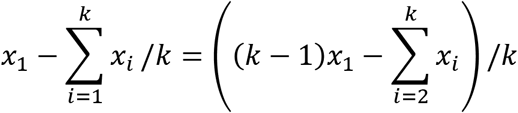

Fourth, define C_1_ = (b_i+1_)^n-k^/k(n-k)! These four steps allow Q_ik_ to be re-written as

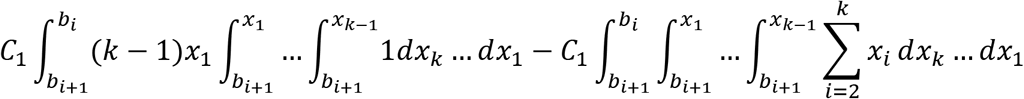

Fifth,

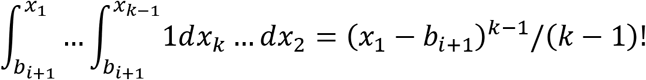

Sixth,

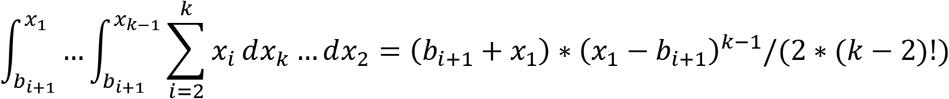

Therefore,

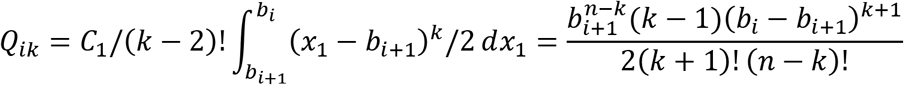

**2.3**: deriving and simplifying TD_i_

Defining TD_i_ as 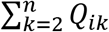 leads to the expression given in TD_i1_, expansion of the binomial exponential reveals DL_i2_, and expansion of the double summation followed by the grouping of terms by exponent of b_i_ reveals DL_i3_.

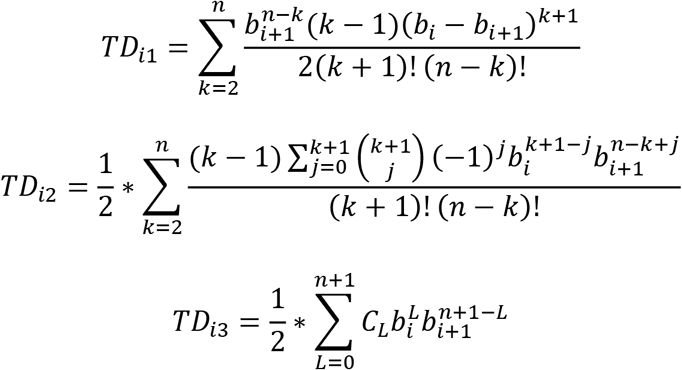

In TD_i3_, the coefficients C_L_ can be derived from noting that in TD_i2_, for all powers of b_i_, there is a one-to-one correspondence between the outer sum’s k values and the inner sum’s j values. Thus,

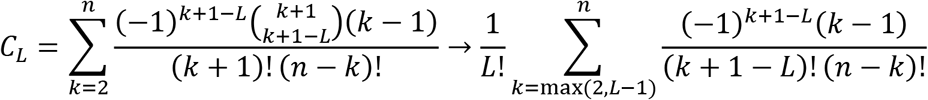

The rightmost expression is the consequence of observing that k+1-L is potentially negative for certain combinations of (k,L), and that in DL_i2_ there is no 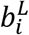 term in the inner sum while L < k+2. This formulation yields

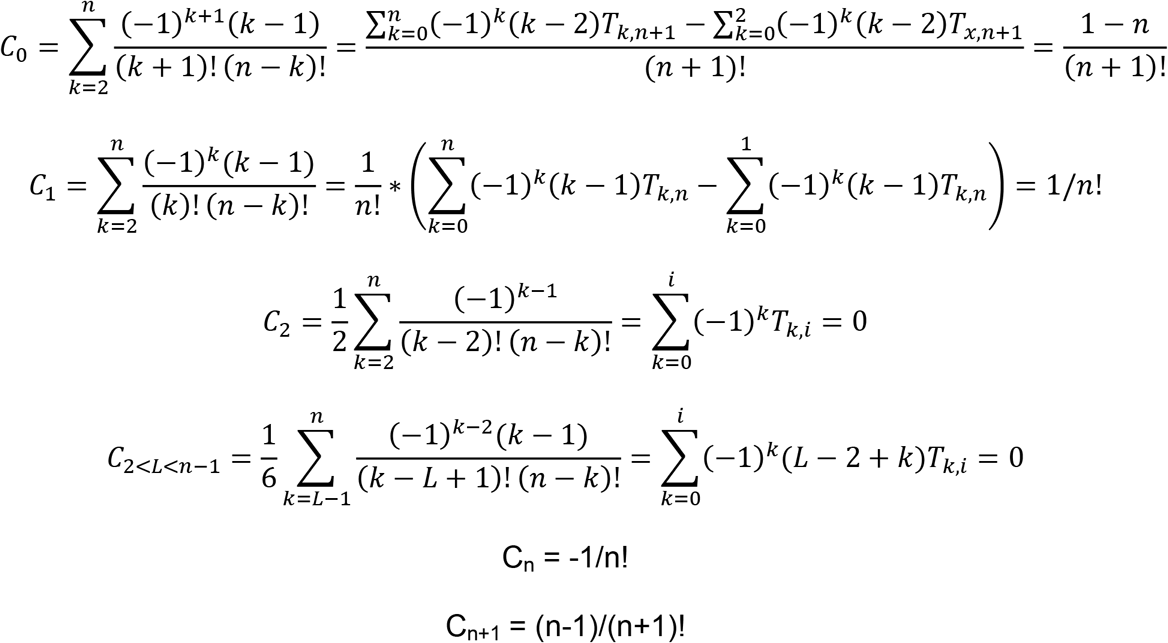

Combining all of the terms derived yields

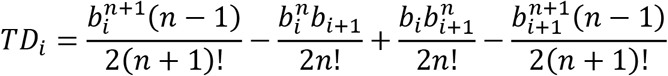

In the above expressions, 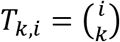 and represents the kth term of the ith row of Pascal’s triangle. In Part 2.3.1 it is shown that certain sums over T_k,i_ terms equal 0.

**2.3.1**: 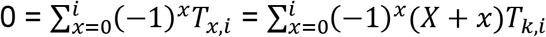 (X is an arbitrary integer)

A well-known feature of Pascal’s triangle is that T_k,i_ = T_k-1,i-1_ + T_k,i-1_ (T_-1,i_ = T_i+1,i_ = 0). Therefore, the T_k,i_ terms in the sum 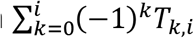 can be replaced with T_k,i-1_ terms; additionally decomposing the sum itself into its positive and negative components yields

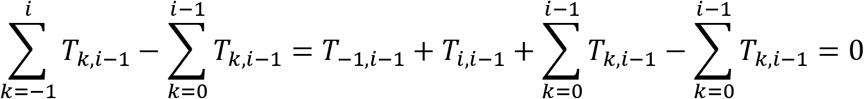

It is also true that T_k,i_ = T_k-2,i-2_ + 2*T_k-1,i-2_ + T_k,i-2_. Proposing that 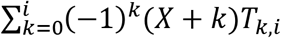 can be expressed as 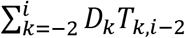 the coefficients D_k_ can be derived by noting first that individual T_k,i-2_ terms only contribute to T_n,i_ for n∈[k:k+2]; within [k:k+2], observe that

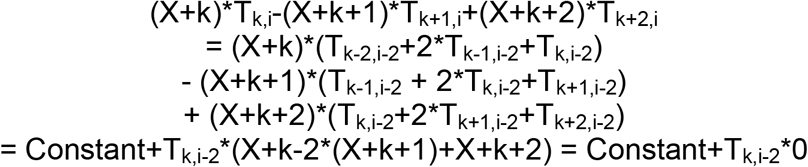

Since the individual D_k_ = 0, 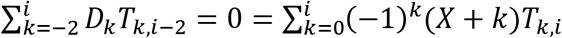

**2.4**: Expression for dTD/db_i_

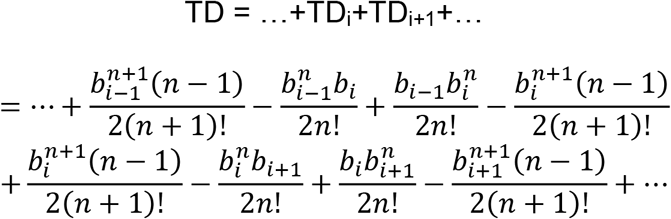

Setting dTD/db_i_ = 0 and solving yields

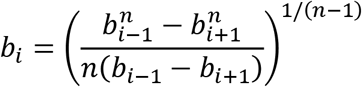

Finally, rewriting TD and U as shown below reveals that they vary in opposite directions with respect to b_i-1_(b_i_)^n^-b_i_(b_i-1_)^n^. So, the b_i_ that maximize utility will minimize distortion.

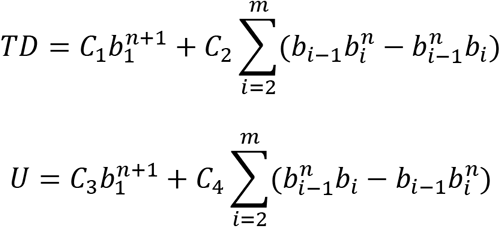

**Part 3**: proof that, for two-item decisions on the attribute-utility landscape in Figure 1b1, the perceptual boundary that optimizes utility also optimizes distortion

In this decision environment, setting a > 1, utility as a function of attribute value x is

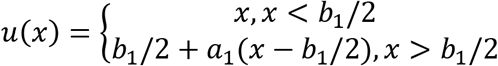

From Part 2.1, for a 1-bit interface, U = p(P_1_)E(P_1_)+p(P_2_)E(P_2_). Also, assuming (reasonably) that b_2_ > b_1_/2,

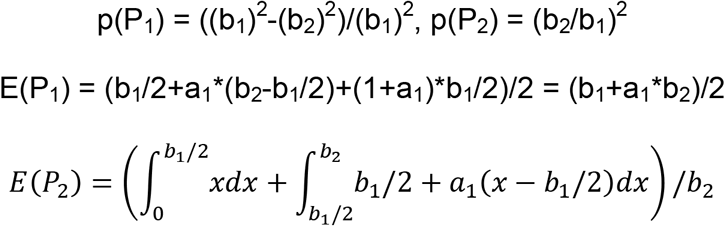

Setting dU/db_2_ = 0 and solving yields b_2_ = b_1_*(5/8-a^−1^/8).

Considering now the optimization of TD, the present decision environment affords three simplifications to the integrand, 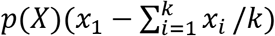. First, p(X) is a constant. Second, 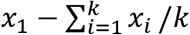 simplifies to (x_1_−x_2_)/2. Third, when x_1_ and x_2_ correspond to different percepts, the integrand is zero. Therefore,

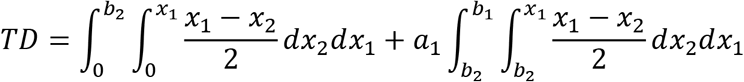

Here, it is illuminating to split the first double integral into the following components:

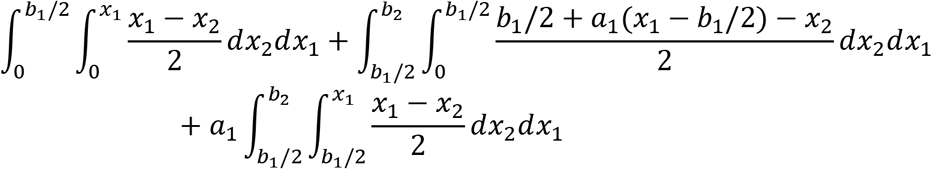

Setting dTD/db_2_ = 0 and solving yields b_2_ = b_1_*(5/8−1/(8*a_1_)).

**Part 4**: proof that, for two-item decisions on the attribute-utility landscape in Figure 1b2, the perceptual boundary that optimizes utility also optimizes distortion

In this decision environment, with a_2_∈[0,1/b_1_], the item PDF as a function of attribute value x, and the resulting utility function, are given below:

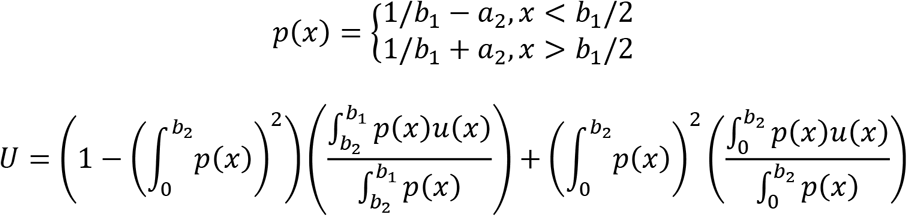

U reduces to a more familiar form when a_2_ = 1. For a_2_∈[0,1/b_1_], 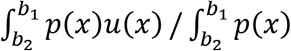 simplifies to (b_1_+b_2_)/2 because p(x) is constant in [b_2_,b_1_]. Also, for specificity’s sake:

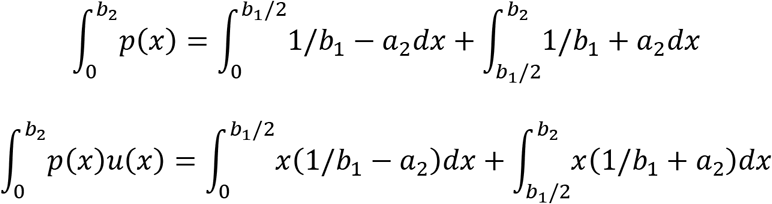

Setting dU/db_2_ = 0 and solving yields b_2_ = b_1_*(1/2+a_2_*b_1_/4), which is a linear function of a_2_ with range [b_1_/2,3*b_1_/4].

Meanwhile, letting p_1_(x_1_,x_2_) = (1/b_1_−a_2_)^2^, p_2_(x_1_,x_2_) = (1/b_1_−a_2_)*(1/b_1_+a_2_), and p_3_(x_1_,x_2_) = (1/b_1_+a_2_)^2^,

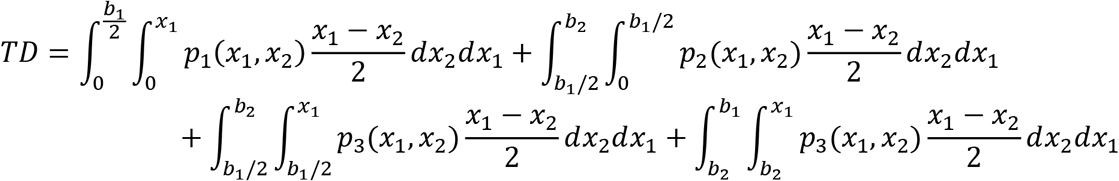

Setting dTD/db_2_ = 0 and solving yields b_2_ = b_1_*(1/2+a_2_*b_1_/4).

**Part 5**: demonstration that the bit-attribute assignments that maximize utility are the same as those that minimize distortion (2 bits, 2 independent and independently-perceived attributes, 2 item choices, uniform item probabilities over attribute values)

**5.1**: definition of problem and notation

The ranges of attributes 1 and 2 are [0,b_1,1_] and [0,b_2,1_], respectively. If item utilities are the sum of attribute values, utility and distortion are abbreviated U_s_ and D_s_, respectively. Alternatively, if utility is situationally-determined, depending on attributes 1 and 2 with probabilities p and 1-p (respectively), I use the abbreviations U_p_ and D_p_. These abbreviations are in turn superscripted according to the attributes in which perceptual bits are invested.

Thus, if both bits are invested in differentiation along attribute 1,

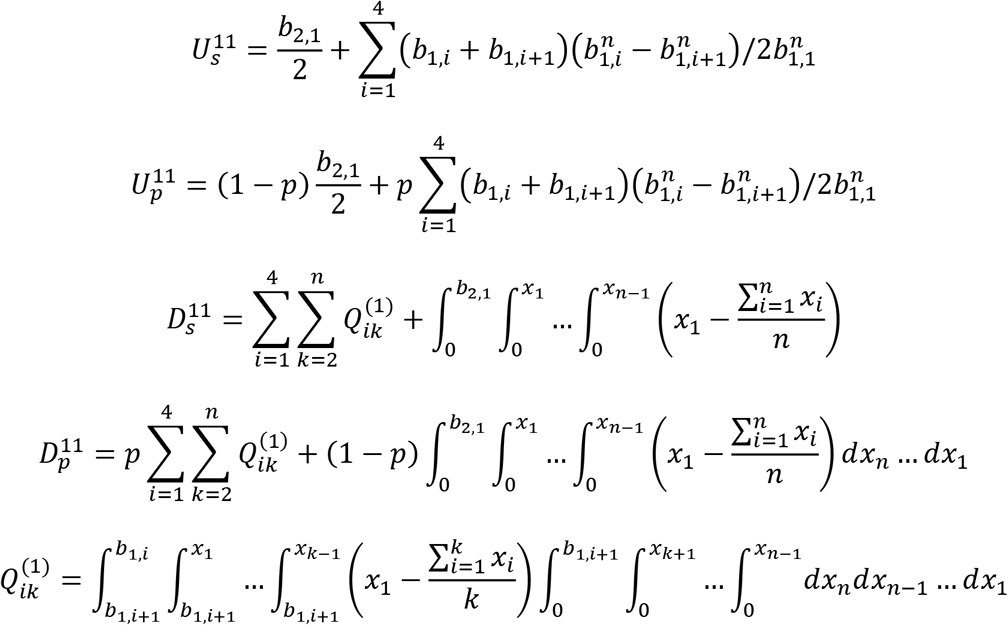

If one bit is invested in differentiation along each attribute,

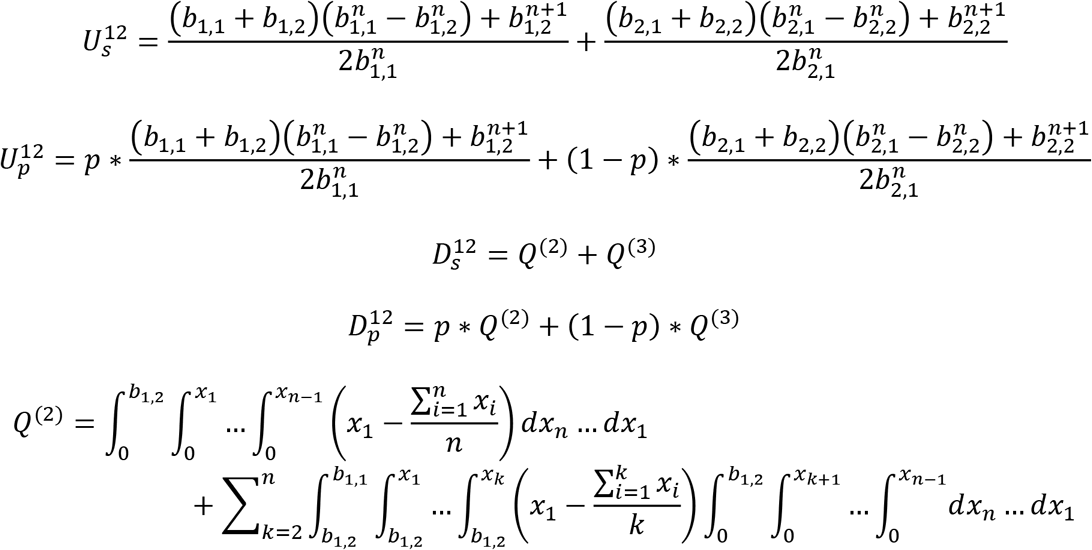

To obtain Q^(3)^ from Q^(2)^, respectively replace b_1,1_ and b_1,2_ with b_2,1_ and b_2,2_.

**5.2**: along attribute a, for a particular n and boundaries b_i_, U_a_ = C_1_*b_a,1_ and D_a_ = C_2_*b_a,1_; so, if b_a,1_ is scaled by f, so are U_a_ and D_a_.

First, from Parts 2.1 and 2.4, 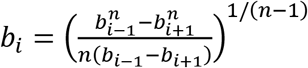. If b_1_ is scaled by f, and all other b_i_ are also scaled by f, then this formula still holds true:

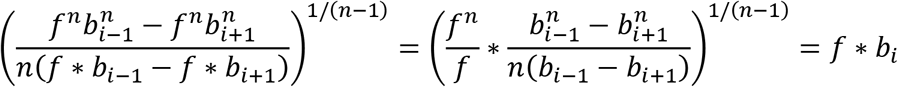

Second, from Part 2.1, 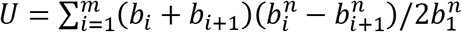; scalling all b_i_ by f gives

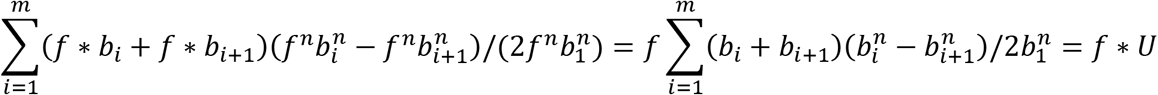

Third, from Parts 1 and 2.4,

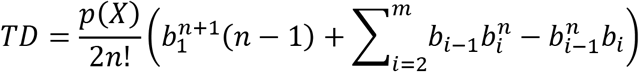

Assuming uniform item probability over each attribute’s span, p(X) = 1/(b_a,1_)^n^. Thus, substituting f*b_a,1_ for b_a,1_,

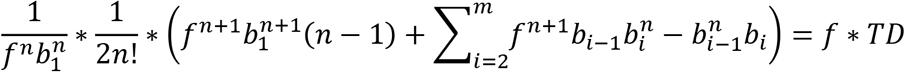

**5.3**: for all U^11^ and D^11^ vs. U^12^ and D^12^, the optimal bit-attribute allocations are analogous

Knowing that utilities and distortions are linear functions of b_a,1_ allows several simplifications to be made to the U^xx^ and D^xx^ expressed in Part 5.1. The simplifications make use of the following expressions (which correspond to n = 2; when β > 1, closed-form expressions are not obtainable for n > 2; see Part 5.3.1):

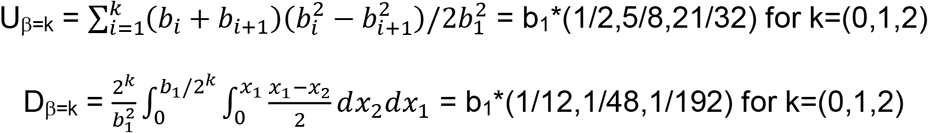

The above solutions use the result from Part 2.4 to determine that b_2_ = b_1_/2 when β = 1, and that (b_2_,b_3_,b_4_) = (.75b_1_,.5b_1_,.25b_1_) when β = 2. For further simplicity, b_1,1_ = 1 and b_2,1_/b_1,1_ = R; now, the U^xx^ and D^xx^ become:

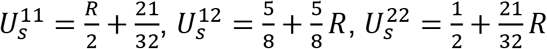

Thus, 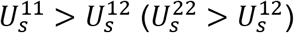 when R < .25(R > 4)

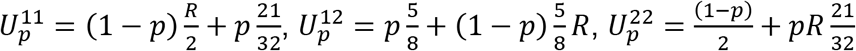

Thus, 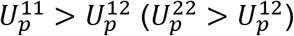 when R*(1-p)/p < .25 (R*(1-p)/p > 4)

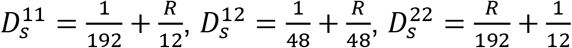

Thus, 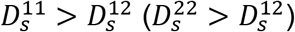 when R < .25(R > 4)

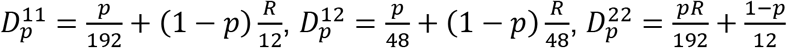

Thus, 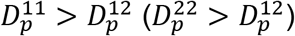 when R*(1-p)/p < .25(R*(1-p)/p > 4)

The bit allocation optimality transitions are equivalent for U_s_ vs. D_s_, and for U_p_ vs. D_p_.

**5.3.1**: derivation of b_i_, from formula in Part 2.4

For β = 0 and β = 1, the derivations are trivial. For β = 2, b_4_ = b_3_*n^−1/(n−1)^, which leads to 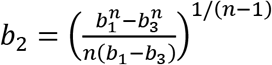, which leads to an expression for b_3_ that is intractable in general but is exactly solvable when n = 2.

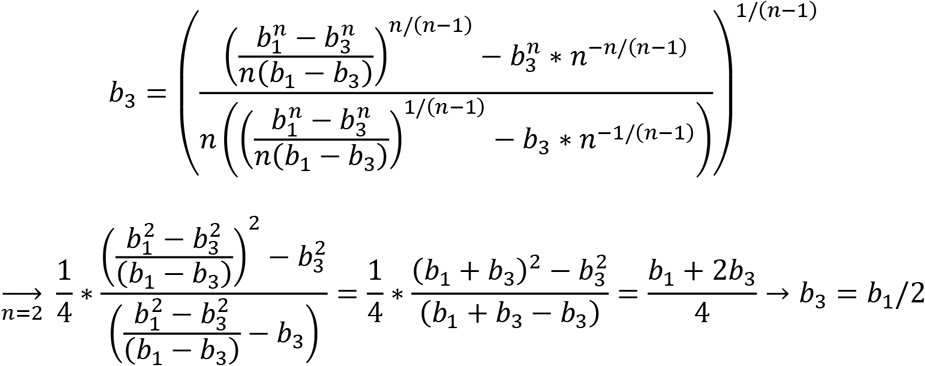

**Part 6**: proof that the percept boundary placement that maximizes evolutionary utility differs from those which either maximize percept-item mutual information or minimize utility estimation error

**6.1**: utility

From Part 2.1, the 1-bit interface is optimized when b_2_=b_1_/n^1/(n-1)^.

6.2: mutual information

If the items spanning 0:b_1_ are chosen at random, their differential entropy (H) is 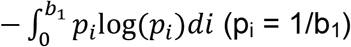.

In the case of the one-bit interface, the conditional differential entropy (H_c_), conditioned on the low- and high-utility percepts (labeled L and H, respectively), equals

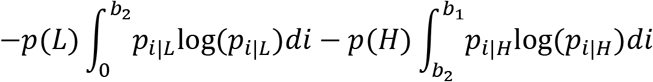

Here, p_i|L_ = 1/b_2_ and p_i|H_ = 1/(b_1_−b_2_).

Mutual information (MI) = H−H_c_, and this holds true whether one is considering the mutual information between percepts and items perceived (MI_P_) or between percepts and items chosen (MI_C_); it should be specified, however, that in the MI_P_ (MI_C_) case, p(L) and p(H) are respectively equal to b_2_/b_1_ and 1-b_2_/b_1_ ((b_2_/b_1_)^n^ and ((b_1_)^n^-(b_2_)^n^)/(b_1_)^n^).

It is clear by inspection that maximizing MI_P_ with respect to b_2_ is not equivalent to maximizing utility because only the latter depends on n.

Deviation between dMI_C_ and utility is demonstrated in Part 6.4.

**6.3**: estimation error

The following expression serves as a generalized mean error function:

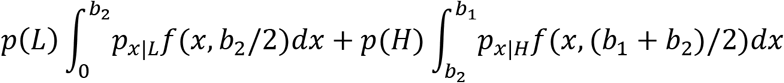

In the absolute and squared error cases, f(a,b) = |a-b| and (a-b)^2^, respectively.

Analogously with Part 6.2, p_x|L_ = 1/b_2_ and p_x|H_ = 1/(b_1_-b_2_); furthermore, when considering the utility estimate error of items perceived (chosen), E_P_ (E_C_), the values of p(L) and p(H) are equivalent to those for MI_P_ (MI_C_). It is therefore still clear by inspection that the maxima of utility and E_P_ are not equivalent.

In Part 6.4 it is confirmed that the maxima of utility and E_C_ are not equivalent either.

**6.4**: further study of MI_C_ and E_C_

Setting dMI_C_/db_2_ = 0 and simplifying yields

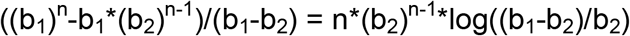

This does not afford an analytic solution for all n; it does not reduce to b_2_ = b_1_/n^1/(n-1)^.

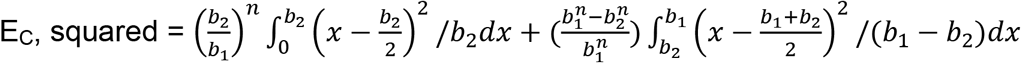

Setting dE_C_/db_2_ = 0 and simplifying yields

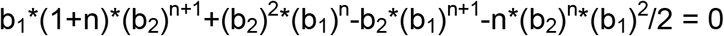

This does not afford an analytic solution for all n; it does not reduce to b_2_ = b_1_/n^1/(n-1)^. (Illustratively, for the case n = 2, it reduces to b_2_ = b_1_/3^1/2^.)

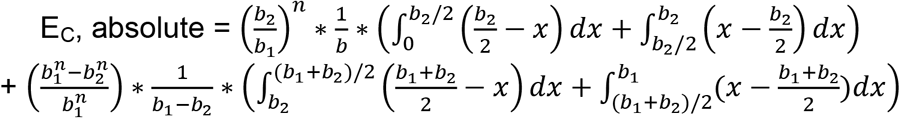

Setting dE_C_/db_2_ = 0 and simplifying yields 2*(n+1)*(b_2_)^n+1^-n*b_1_*(b_2_)^n^-b_2_*(b_1_)^n^ = 0.

This does not afford an analytic solution for all n; it does not reduce to b_2_ = b_1_/n^1/(n-1)^. (Illustratively, for the case n = 2, it reduces to b_2_ = (1+7^1/2^)*b_1_/6.)

**Part 7**: proof that the percept boundaries that minimize estimation error across all percepts correspond to an optimum of the distortion loss and utility objectives when 1) attribute probabilities are uniform 2) max-utility attribute values are uniformly probable 3) utility varies linearly with attribute value 4) choices are binary in a linear attribute space (spanning [0,b_1_]) OR the attribute space is radial (spanning [−π,π])

Let point p(p∈[0,b_1_]) emerge as the randomly-determined peak utility attribute value; denote the corresponding distortion and utility functions as D_p_ and U_p_, respectively.

The objective function corresponding to the interface’s overall utility is 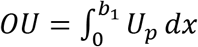. To optimize the boundary placement of a 1-bit interface, set dOU/db_2_ = 0 and solve.

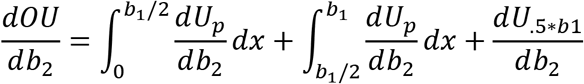

Estimation error is minimized when b_2_ = .5*b_1_; fortuitously, at this value of b_2_, the integral terms are additive inverses and dU._5*b1_/db_2_ = 0. Moreover, this result is independent of n (the number of item choices presented).

To construct an optimal 2-bit interface, it is instructive to assume that b_1_/2 is the optimal placement for b_3_ and to define OU_1_ and OU_2_ such that U_p_ ∈ OU_1_ (OU_2_) when p ∈ [0,b_1_/2] (p ∈ [b_1_/2,b_1_]). Within the interval [b_3_,b_1_],

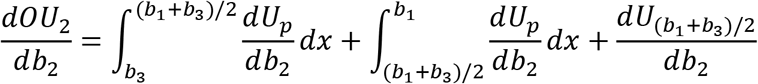

By analogy with the 1-bit interface, the optimal placement of b_2_ with respect to OU_2_ is midway between b_1_ and b_3_. This analogy extends to β-bit interfaces as follows: the optimal placement of b_i_ with respect to OU_i_, U_p_ ∈ OU_i_ when p ∈ [b_i+1_,b_i−1_], is (b_i−1_+b_i+1_)/2.

Meanwhile, within the interval [b_3_,b_1_], all U_p_ ∈ OU_1_ reduce to c-x (c constant); the division of these linear utility gradients is therefore equivalent and as described in Part 2. Thus, the placement of b_2_ at (b_3_+b_1_)/2 is optimal uniquely when n = 2. This result extends to β-bit interfaces as follows: with n = 2, the optimal placement of b_i_ with respect to OU_x_, U_p_ ∈ OU_x_ when p ¬∈ [b_i+1_,b_i−1_], is (b_i−1_+b_i+1_)/2.

All of these constructions generalize to the radial attribute space, with one exception. Consider, in a linear attribute space, the placement of b_i_ within [b_i+1_,b_i-1_]. If b_i−1_ ≤ b_1_/2 (an analogous argument can be made if b_i+1_ ≥ b_1_/2), then define: b_m+1_ = 0, dh = b_i+1_+b_i−1_, U_p_ ∈ OU_1_ (OU_2_, OU_3_, OU_4_) when p ∈ [b_m+1_,b_i+1_] ([b_i+1_,b_i−1_], [b_M_,b_dh_], [b_dh_,b_1_]). Furthermore, assume uniform spacing of all existing boundaries prior to the insertion of b_i_. Under these conditions, for any n, at b_i_ = (b_i-1_+b_i+1_)/2, dOU_1_/db_i_ = −dOU_3_/db_i_ and dOU_2_/db_i_ = 0; however, dOU_4_/db_i_ = 0 only if n = 2. In a radial attribute space, meanwhile, rotational symmetry allows boundary labels to be rotated arbitrarily; so, let the boundary placement problem be framed as the placement of bo between b_1_ and b_m_, b_i_ (i = 1:m) spaced evenly. It is optimal to place bo midway between b_1_ and b_m_; ∀ k ∈ [0,b_m_−b_1_], dU_1+k_/db_0_ = −dU_m-k_/db_0_.

Finally, the above expressions continue to hold true if D_p_ is substituted for U_p_, and so do the corresponding conclusions.

**Part 8**: proof that in a radial attribute space, when 1) utility varies linearly with attribute value 2) max-utility attribute values are uniformly probable 3) attribute probabilities belong to a particular but not special non-uniform distribution, utility is optimized by efficient coding of the item probability distribution specifically when decisions are binary, i.e. when n = 2, and that as n increases, utility optimization converges to the efficient coding of the attribute space. (Here, efficient coding is defined as the minimization of attribute value estimation error.)

**8.1**: n = 2

The expression for the total estimation error (TE) can be constructed as follows. First, consider that ∀ x ∈ [−π,π], expected error E(x) can be computed, and the E(x) can then be summed; symbolically, 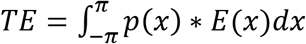. Next, denoting x’s assigned percept P_x_, E(x) is the product of |x-E(P_x_)| times the probability that errors of this magnitude will be realized. The latter of these multiplied quantities can be written as 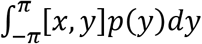, where [x,y] = 1 (0) if x and y are (not) assigned to the same percept. Combining these ideas yields

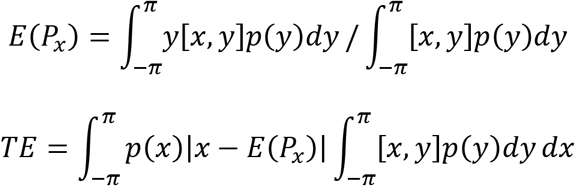

Next, note that 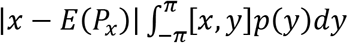 expands as follows:

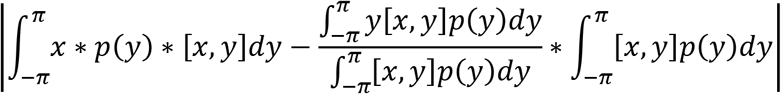

Simplifying the above expression yields an integrand over y of |x-y|p(y)[x,y]; if p(x) and p(y) are further combined into p(x,y), then

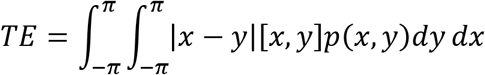

An expression for utility can be constructed similarly. First, ∀ p ∈ [−π,π], expected utility U_p_ can be computed, and the U_p_ can then be summed; symbolically, 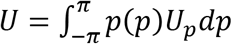. Second, the peak value can be denoted as M. Third, denoting the utility obtained from the sampling of item pair (x,y) as U_pxy_, 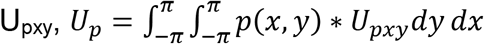. Finally, if x and y are assigned to different percepts, then U_pxy_ = min(|p-x|,|p-y|); otherwise, U_pxy_ is the average of min(|p-x|,|p-y|) and max(|p-x|,|p-y|). Combining these ideas yields

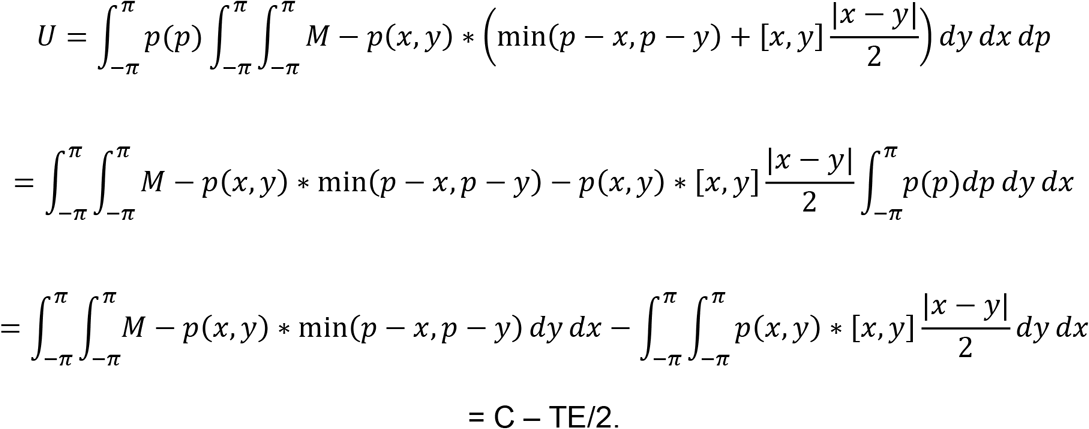

In the above expressions, note that 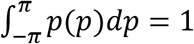 and that Cisa constant. This simplification clarifies that TE and U depend equally and oppositely on the values of [x,y], which are determined by the item-percept assignment; thus, they are optimized by the same assignment.

**8.2**: large n

For every potential peak utility attribute value p, let p(P_p_) be the probability that, from a set of n sampled items, at least 1 belongs to the same percept assignment group as p. Also, let E(P_p_) E(¬P_p_) respectively be the expected values derived when at least one and when none of the sampled items belong to P_p_. Then,

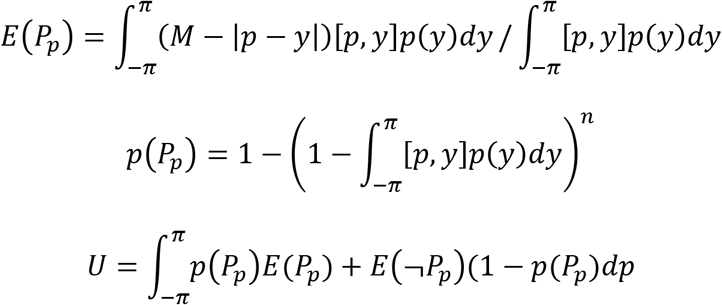

E(P_p_) is unaffected by n, while as n grows, p(P_p_) converges to 1. This convergence allows U to be approximated as

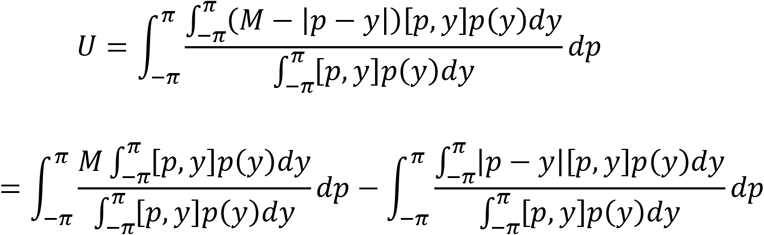

The first of these integrals is a constant, while the minimization of the second maximizes U. In fact, for optimization purposes, the quotient of integrals (Q) can be considered in isolation.

To illustrate the value of efficiently encoding the attribute space, I assume β = 2 and

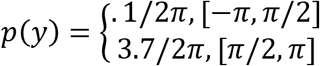

Q is straightforwardly minimized by placing boundaries at −π/2, 0, π/2, and π; this boundary placement diverges obviously from the efficient coding of p(y), which would make distinctions within the interval [π/2,π].

